# Magnesium increases homoeologous crossover frequency in *ZIP4* (*Ph1*) mutant wheat-wild relative hybrids

**DOI:** 10.1101/278341

**Authors:** María-Dolores Rey, Azahara C. Martín, Mark Smedley, Sadiye Hayta, Wendy Harwood, Peter Shaw, Graham Moore

**Author notes:** Corresponding author: María-Dolores Rey.

## Abstract

Wild relatives provide an important source of useful traits in wheat breeding. Wheat and wild relative hybrids have been widely used in breeding programs to introduce such traits into wheat. However, successful introgression is limited by the low frequency of homoeologous crossover (CO) between wheat and wild relative chromosomes. Hybrids between wheat carrying a 70Mb deletion on chromosome 5B (*ph1b*) and wild relatives, have been exploited to increase the level of homoeologous CO, allowing chromosome exchange between their chromosomes. In *ph1b*-rye hybrids, CO number increases from a mean of 1 CO to 7 COs per cell. CO number can be further increased up to a mean of 12 COs per cell in these *ph1b* hybrids by treating the plants with Hoagland solution. More recently, it was shown that the major meiotic crossover gene *ZIP4* on chromosome 5B (*TaZIP4-B2*) within the 70Mb deletion, was responsible for the restriction of homoeologous COs in wheat-wild relative hybrids, confirming the *ph1b* phenotype as a complete *Tazip4-B2* deletion mutant (*Tazip4-B2 ph1b)*. In this study, we have identified the particular Hoagland solution constituent responsible for the increased chiasma frequency in *Tazip4-B2 ph1b* mutant-rye hybrids and extended the analysis to *Tazip4-B2* TILLING and CRISPR mutant-*Ae variabilis* hybrids. Chiasma frequency at meiotic metaphase I, in the absence of each Hoagland solution macronutrient (NH_4_ H_2_PO_4_, KNO_3_, Ca (NO_3_)2·4H_2_O or Mg SO_4_·7H_2_O) was analysed. A significant decrease in homoeologous CO frequency was observed when the Mg^2+^ ion was absent. A significant increase of homoeologous CO frequency was observed in all analysed hybrids, when plants were irrigated with a 1mM Mg^2+^ solution. These observations suggest a role for magnesium supplementation in improving the success of genetic material introgression from wild relatives into wheat.

## Introduction

Despite possessing related ancestral genomes (genome AABBDD), bread wheat behaves as a diploid during meiosis. Deletion of chromosome 5B in tetraploid and hexaploid wheat results in a level of incorrect chromosome pairing and exchange, visualised as a low level of multivalents at metaphase I, and homoeologous crossovers (COs) between related chromosomes in wheat-wild relative hybrids (Riley and Chapman, 1958; Sears and Okamoto, 1958). From these observations, it was proposed that chromosome 5B carries a locus termed *Pairing homoeologous 1* (*Ph1*), which evolved on wheat’s polyploidisation and restricted chromosome pairing and COs to true homologues (Riley and Chapman, 1958). A hexaploid wheat cv. Chinese Spring (CS) line carrying a 70Mb deletion on the long arm of chromosome 5B (*ph1b*) has been exploited over the last 40 years to allow exchange between wild relative and wheat chromosomes. Recently, it was shown that on wheat’s polyploidisation, a segment of 3B carrying the major crossover gene *ZIP4* and a block of heterochromatin, duplicated and inserted between two *CDK2*-like genes within a cluster of *CDK2-like* and methyl-transferase genes (Griffiths et al., 2006; Al-Kaff et al., 2008; Martín et al., 2014, 2017). Using exploitation of TILLING mutants, it was shown that the duplicated *ZIP4* gene (*TaZIP4-B2*) within this cluster, both promotes homologous CO and restricts homoeologous CO (Rey et al., 2017). Therefore, *TaZIP4-B2* within the 70Mb *ph1b* deletion region is responsible for the effect on homoeologous CO in wheat-wild relative hybrids, and as such the *ph1b* line can be described as a complete-deletion (or complete loss-of-function) mutant of *Tazip4-B2 (Tazip4-B2 ph1b* mutant). In terms of the effect on chromosome synapsis/pairing, cell biological studies reveal that the *ph1b* deletion in wheat has little effect, with most synapsis occurring during clustering of the telomeres as a bouquet. Furthermore, in wheat-wild relative hybrids, which only possess homoeologues, the *ph1b* deletion also has little effect on the level of synapsis, except that most pairing occurs after dispersal of the telomere bouquet. In wheat itself, a few chromosomes also undergo delayed pairing until after dispersal of the bouquet, with the subsequent incorrect pairing leading to the low level of multivalents observed at metaphase I (Martín et al., 2014, 2017).

For the last 40 years, the wheat CS *ph1b* deletion line has been exploited in crosses with wild relatives to allow exchange between chromosomes at meiosis. As indicated previously, in these hybrids, the extent of chromosome synapsis is similar whether the line carries the *ph1b* deletion or not. Moreover, on the synapsed chromosomes, similar numbers of MLH1 sites (normally a marker for CO), are observed (Martín et al., 2014). However significant site CO frequency is only observed in those hybrids carrying the *ph1b* deletion. However even in this case, the frequency of resulting COs still does not reflect the number of available MLH1 sites (Martín et al., 2014). This implies that there is potential for increased processing of MLH1 sites into COs. Fortuitously, it has been observed that a nutrient solution (Hoagland’s solution) added to the soil when *Tazip4-B2 ph1b* mutant-rye hybrids are growing resulted in increased CO frequency, although it was not known which nutrient component was responsible for the effect.

Mineral elements are essential nutrients for plants to complete their life cycle. They are classified into macro and micronutrients, which are required in relatively large and small amounts, respectively (Hoagland and Arnon, 1950). The importance of each of these macronutrients has been reported in numerous physiological processes, such as plant growth, cell division, and metabolism (Huber, 1980; Maathius, 2009). However, limited studies have been performed as to their effect on meiosis. Early studies have previously reported that alterations of external factors, such as temperature, or nutrient composition, can produce profound effects on chiasma frequency (Grant, 1952; Wilson, 1959; Law, 1963; Bennet and Ress, 1970; Fedak, 1973).

The main objective of the present study was to determine whether a specific macronutrient present in the Hoagland solution was responsible for the observed increased homoeologous CO frequency in *Tazip4-B2 ph1b* mutant-rye hybrids described in Martín et al., (2017). We also analysed whether this macronutrient increased homoeologous CO frequency in each of the *Tazip4-B2 ph1b* (complete deletion), TILLING (point mutation) and CRISPR (partial deletion) mutant-*Ae. variabilis* hybrids.

## Materials and methods

### Plant material

Plant material used in this study included: *Triticum aestivum* (2n = 6x = 42; AABBDD) cv. Chinese Spring *Tazip4-B2*-*ph1b* mutant line (Sears, 1977); *Triticum aestivum* cv. Chinese Spring-rye hybrids crosses between the *Tazip4-B2*-*ph1b* mutant line hexaploid wheat and rye (*Secale cereal* cv. Petkus (2n = 2x= 14; RR)); *Triticum aestivum* cv. Chinese Spring-*Aegilops variabilis* hybrids – crosses between *Tazip4-B2*-*ph1b* mutant and *Ae. variabilis* Eig. (2n = 4x = 28; UUS^v^S^v^); *Triticum aestivum* cv. Cadenza-*Ae. variabilis* hybrids - crosses between Cad1691 and Cad0348, *Tazip4-B2* TILLING mutants and *Ae. variabilis* (Krasileva et al., 2016; Rey et al., 2017)); and *Triticum aestivum* cv. Fielder-*Ae. variabilis* hybrids crosses between *Tazip4-B2* CRISPR mutant and *Ae. variabilis* (see Production of *TaZIP4-B2* knock-out using RNA-guided Cas9, Materials and Methods).

### Nutrient solution treatments

The total number of plants used in this work is described in Table S1. All seedlings were vernalised for 3 weeks at 7°C under a photoperiod of 16h light/8h dark, and then transferred to a controlled environmental room until meiosis (approximately 2 months later for all genotypes used in this study). The growth conditions were 16h/8h, light/dark photoperiod at 20°C day and 15°C night, with 70% humidity. At least two weeks before meiosis, irrigation of plants with a Hoagland solution (100 mL per plant) was commenced following the method previously described in Martín et al., (2017). Briefly, plants were irrigated once a week with a Hoagland solution (100mL) from the stem elongation stage of the vegetative stage (stage 7-8, Feeke’s scale). The composition of the Hoagland solution was: (*macronutrients*) KNO_3_ (12 mM), Ca (NO_3_)2·4H_2_O (4 mM), NH4 H_2_PO_4_ (2 mM), Mg SO_4_·7H_2_O (1 mM); and (*micronutrients*) NaFe-EDTA (60 mM), KCl (50 μM), H_3_BO_3_ (25 μM), Mn SO_4_·H_2_O (2 μM), Zn SO_4_ (4 μM), Cu SO_4_·5H_2_O (0.5 μM), H_2_MoO_4_ (0.5 μM). Four treatments were carried out to analyse the effect of the absence of NH_4_ H_2_PO_4_, KNO_3_, Ca (NO_3_)2·4H_2_O or Mg SO_4_·7H_2_O from the Hoagland solution on homoeologous CO frequency in *Tazip4-B2*-*ph1b* mutant-rye hybrids. For each treatment, a different Hoagland solution was prepared in the absence of each macronutrient (NH_4_ H_2_PO_4_, KNO_3_, Ca (NO_3_)2·4H_2_O or Mg SO_4_·7H_2_O). Moreover, the effect of the presence of only Mg SO_4_·7H_2_O (Mg SO_4_·7H_2_O is designed as Mg^2+^ in the manuscript) in only water rather than in Hoagland solution on CO frequency was also analysed in *Tazip4-B2 ph1b* mutant-rye hybrids in comparison to the effects of Hoagland solution. Also, two different concentrations of Mg^2+^ (1mM and 2mM of Mg^2+^ in water) were used to assess the homoeologous CO frequency in *Tazip4-B2*-*ph1b* mutant-rye hybrids. The treatment with either Mg^2+^ in water alone or Hoagland solution in *Tazip4-B2 ph1b* mutant*-Ae. variabilis*, and *Tazip4-B2* TILLING and CRISPR mutant-*Ae. variabilis* hybrids was also assessed.

Assessment of the addition of Mg^2+^ in water alone on homoeologous CO frequency was also made on non-irrigated plants, by injecting into *Tazip4-B2 ph1b* mutant-rye hybrids tillers a solution containing 1mM Mg^2+^ in water (0.5 mL per spike) just above every spike (three spikes with water alone and three spikes with Mg^2+^ in water). All spikes were analysed 24-48h after the injection.

### Feulgen-stained analysis

After either irrigating with Hoagland or Mg^2+^ solution, or injecting the Mg^2+^ solution, tillers were harvested when the flag leaf was completely emerged, and only anthers at meiotic metaphase I were collected and fixed in 100% ethanol/acetic acid 3:1 (v/v). The anthers used in this study were taken from spikelets in the lower half of the spike. From each spikelet, the 2 largest florets (on opposing sides of the floret) were used. From each dissected floret, one of the three synchronised anthers was squashed in 45% acetic acid/distilled water (v/v) and the meiocytes assessed for being at meiotic metaphase I by observation under a phase contrast microscope (LEICA DM2000 microscope (LeicaMicrosystems, http://www.leica-microsystems.com/)). The two remaining anthers were left then fixed in 100% ethanol/acetic acid 3:1 (v/v) for cytological analysis of meiocytes. The anthers were incubated in ethanol/acetic acid at 4°C for at least 24h. Cytological analysis of meiocytes at metaphase I was performed using Feulgen reagent as previously described in Sharma and Sharma, (2014). Metaphase I meiocytes were observed under a phase contrast microscope equipped with a Leica DFC450 camera and controlled by LAS v4.4 system software (Leica Biosystems, Wetzlar, Germany). The digital images were used to determine the meiotic configurations of the meiocytes by counting the number of univalents, rod (1 chiasma) and ring (2 chiasmata) bivalents and multivalents (trivalents (1-2 chiasmata), tetravalents (3 chiasmata) and pentavalents (4 chiasmata)). Two different methods depending on the number of chiasma (single or double chiasmata) were used to calculate chiasma frequency per meiocyte (see Figure S1 for examples of the scored structures). Images were processed using Adobe Photoshop CS5 (Adobe Systems Incorporated, US) extended version 12.0 ×64.

### Production of *TaZIP4-B2* CRISPR mutants using RNA-guided Cas9

Three single guide RNAs (sgRNA) were designed manually to specifically target *TaZIP4-B2*. These guides were in the limited regions where there was sufficient variation between *ZIP4* on 5BL and homoeologous group 3 chromosomes (Figure 3). The specific guides were: Guide 4: 5’GATGAGCGACGCATCCTGCT3’, Guide 11: 5’GATGCGTCGCTCATCCTCCG3’ and Guide 12: 5’GAAGAAGGATGCGGCCTTGA3’ (Figure 3). Two constructs were assembled using standard Golden Gate assembly (Werner et al., 2012) with each construct containing the Hygromycin resistance gene under the control of a rice Actin1 promoter, Cas9 under the control of the rice ubiquitin promoter and two of the sgRNAs each under the control of a wheat U6 promoter (Figure 3). Construct 1 contained guides 4 and 12 and construct 2 contained guides 11 and 12. To produce each gRNA, a PCR reaction was performed using Phusion High-Fidelity Polymerase (Thermo Scientific M0530S) with a forward primer containing the gRNA sequence, and a standard reverse primer 5´TGTGGTCTCAAGCGTAATGCCAACTTTGTAC3´ using the plasmid pICSL70001∷U6p∷gRNA (Addgene plasmid 46966) as template. Each gRNA was cloned individually into the level 1 vectors pICH47751 (gRNA4 & 11) and pICH47761 (gRNA12). Level 1 construct pICH47802-RActpro∷Hpt∷NosT (selection maker), pICH47742-RUbipro∷Cas9∷NosT and the gRNAs were then assembled in the binary Level 2 vector pAGM8031 (Addgene 48037) (Figure 3).

The two constructs were introduced to *T. aestivum* cv. Fielder by Agrobacterium-mediated inoculation of immature embryos. 450 immature embryos were inoculated with Agrobacterium strain AGL1 containing each construct. Briefly, after 3 days co-cultivation with Agrobacterium, immature embryos were selected on 15 mg/l hygromycin during callus induction for 2 weeks and 30 mg/l hygromycin for 3 weeks in the dark at 24°C on Murashige and Skoog medium (MS; Murashige and Skoog, 1962) 30 g/l Maltose, 1.0 g/l Casein hydrolysate, 350 mg/l Myo-inositol, 690 mg/l Proline, 1.0 mg/lThiamine HCl (Harwood et al., 2009) supplemented with 2 mg/l Picloram, 0.5 mg/l 2,4-Dichlorophenoxyacetic acid (2,4-D). Regeneration was under low light (140 μmol.m-2.s-1) conditions on MS medium with 0.5 mg/l Zeatin and 2.5 mg/l CuSO45H2O.

Primary transgenic plants (T0) were analysed by PCR across the region of interest. The sequences for the forward and reverse primers used for the screening in T0 were 5’AGTGGTGAATCCATCCCTTG3’ and 5’CCTTCCTCTTCTTGCACTGG3’, respectively (Rey et al., 2017), followed by direct sequencing. The PCR was performed using RedTaq ReadyMix PCR Reaction Mix (Sigma, St. Louis, MO, USA; R2523) according to the manufacturer’s instructions. PCR conditions were: 3 min 95C, 35 cycles of 15s at 95C, 15s at 58C and 30s at 72C. T0 plants with edits in *TaZIP4-B2* were progressed to the T1 generation and 24 T1 seedlings from each original T0 plant were analysed in the same way for the presence of edits.

### Statistical analyses

Statistical analyses were performed using STATISTIX 10.0 software (Analytical Software, Tallahassee, FL, USA). Analysis of variance (ANOVA) in *Tazip4-B2 ph1b* mutant-rye hybrids, *Tazip4-B2* TILLING mutant-*Ae. variabilis* hybrids and *Tazip4-B2* CRISPR mutant-*Ae. variabilis* hybrids was based on a completely randomised design. Several transformations were carried out: tangent (ring bivalents), arcsine (trivalents) and logarithm (double CO) transformations in the analysis of the effect of absence of each macronutrients in homoeologous CO frequency in *Tazip4-B2 ph1b* mutant-rye hybrids; exponential (ring bivalents) transformation in *Tazip4-B2 ph1b* mutant-*Ae. variabilis* hybrids; exponential (rod bivalents, rings bivalents and trivalents) transformation in *Tazip4-B2* TILLING mutant (Cad1691)-*Ae. variabilis* hybrids; and square root (ring bivalents) and exponential (trivalents) transformations in *Tazip4-B2* TILLING mutant (Cad0348)-*Ae. variabilis* hybrids. Means were separated using the Least Significant Difference (LSD) test with a probability level of 0.05. Both *Tazip4-B2* CRISPR mutant lines and *Tazip4-B2* CRISPR mutant-*Ae. variabilis* hybrids were analysed by the Kruskal–Wallis test (nonparametric one-way analysis of variance). Means were separated using the Dunn’s test with a probability level of 0.05.

## Results

### Magnesium increases homoeologous COs in *Tazip4-B2 ph1b* mutant-rye hybrids

The *Tazip4-B2 ph1b* mutant-rye hybrids were obtained by crosses between the hexaploid wheat cv. Chinese Spring *Tazip4-B2 ph1b* mutant and rye. These hybrids were used to analyse which macronutrient (NH_4_ H_2_PO_4_, KNO_3_, Ca (NO_3_)2·4H_2_O or Mg SO_4_·7H_2_O) present within in the Hoagland solution detailed in Martín et al., (2017) could be responsible for the increased CO number observed in the *Tazip4-B2 ph1b* mutant-rye hybrids. To assess the effect of the absence of each macronutrient in homoeologous CO frequency in meiotic metaphase I, we irrigated several *Tazip4-B2 ph1b* mutant-rye hybrids with: 1) Hoagland solution; 2) water alone; 3) Hoagland solution minus KNO_3_; 4) Hoagland solution minus Ca (NO_3_)2·4H_2_O; 5) Hoagland solution minus NH_4_ H_2_PO_4_; 6) Hoagland solution minus MgSO_4_·7H_2_O (MgSO_4_·7H_2_O is designed as Mg^2+^ in the manuscript) (Table 1). The absence of each Hoagland solution macronutrient caused a slight increase in homoeologous CO frequency, except for the treatment lacking Mg^2+^, where a significant decrease in homoeologous CO frequency per meiocyte was observed at meiotic metaphase I in these hybrids (Table 1). No significant differences in CO frequency at metaphase I were observed between hybrids treated with water alone and those treated with the Hoagland solution minus Mg^2+^ (a mean of 7.91 chiasmata for hybrids treated with water alone and 8.09 chiasmata for hybrids treated with Hoagland solution minus Mg^2+^ (Table 1)).

**TABLE 1:** Effect of the absence of each macronutrient in the Hoagland solution on the homoeologous CO frequency of *T. aestivum* cv. Chinese Spring *Tazip4-B2 ph1b* mutant-rye hybrids. Frequencies of univalents, bivalents, trivalents and chiasma frequency (single and double chiasmata) were scored at meiotic metaphase I in *Tazip4-B2 ph1b* mutant-rye hybrids. Values in parenthesis indicate range of variation between cells. P < 0.05 indicates significant differences according to LSD test.

|  | No. of cell<br>examined | Univalents | Rod bivalents | Ring bivalents | Trivalents | Chiasma frequency | Chiasma frequency |
| --- | --- | --- | --- | --- | --- | --- | --- |
|  |  | Mean ± SE<br>(Range) | Mean ± SE<br>(Range) | Mean ± SE<br>(Range) | Mean ± SE<br>(Range) | Mean ± SE<br>(Range) | Mean ± SE<br>(Range) |
|  |  |  |  |  |  | Single Chiasma | Double Chiasmata |
| Hoagland solution | 109 | 12.36 ± 0.21<sup>c</sup><br>(7-18) | 4.39 ± 0.14<sup>a</sup><br>(2-8) | 2.57 ± 0.10<sup>abc</sup><br>(0-5) | 0.58 ± 0.07<sup>a</sup><br>(0-3) | 10.74 ± 0.16<sup>a</sup><br>(7-14) | 12.03 ± 0.22<sup>a</sup><br>(7-18) |
| Water alone | 108 | 15.93 ± 0.16<sup>a</sup><br>(12-20) | 4.03 ± 0.13<sup>ab</sup><br>(1-7) | 1.68 ± 0.09<sup>bc</sup><br>(0-4) | 0.22 ± 0.04<sup>bc</sup><br>(0-1) | 7.91 ± 0.13<sup>d</sup><br>(5-10) | 8.15 ± 0.15<sup>d</sup><br>(5-12) |
| Hoagland solution - NH<sub>4</sub> H<sub>2</sub>PO<sub>4</sub> | 92 | 13.61 ± 0.22<sup>b</sup><br>(8-17) | 3.83 ± 0.14<sup>b</sup><br>(2-8) | 2.78 ± 0.13<sup>a</sup><br>(0-6) | 0.39 ± 0.06<sup>ab</sup><br>(0-2) | 10.25 ± 0.18<sup>b</sup><br>(7-14) | 11.03 ± 0.23<sup>b</sup><br>(7-17) |
| Hoagland solution - KNO<sub>3</sub> | 93 | 13.86 ± 0.22<sup>b</sup><br>(6-18) | 4.34 ± 0.16<sup>a</sup><br>(2-8) | 2.35 ± 0.12<sup>abc</sup><br>(0-5) | 0.25 ± 0.05<sup>c</sup><br>(0-2) | 9.57 ± 0.18<sup>c</sup><br>(7-16) | 9.96 ± 0.19<sup>c</sup><br>(7-16) |
| Hoagland solution - Ca (NO<sub>3</sub>)<sub>2</sub>·4H<sub>2</sub>O | 102 | 13.84 ± 0.20<sup>b</sup><br>(9-18) | 4.14 ± 0.13<sup>ab</sup><br>(1-7) | 2.50 ± 0.09<sup>c</sup><br>(0-4) | 0.29 ± 0.05<sup>bc</sup><br>(0-2) | 9.79 ± 0.15<sup>c</sup><br>(7-14) | 10.12 ± 0.17<sup>c</sup><br>(7-15) |
| Hoagland solution - Mg SO<sub>4</sub>·7H<sub>2</sub>O | 90 | 15.63 ± 0.24<sup>a</sup><br>(10-20) | 4.12 ± 0.16<sup>ab</sup><br>(1-8) | 1.68 ± 0.13<sup>ab</sup><br>(0-4) | 0.26 ± 0.06<sup>bc</sup><br>(0-2) | 8.09 ± 0.20<sup>d</sup><br>(5-12) | 8.62 ± 0.22<sup>d</sup><br>(5-13) |
| *P-value* |  | 0.0000 | 0.0538 | 0.0592 | 0.0516 | 0.0000 | 0.0000 |

Additionally, we scored all meiocytes for the occurrence of double chiasmata in the metaphase I chromosomal configurations (examples highlighted by arrows in Figure S1). When double chiasmata were considered in the chiasma frequency, a mean of 8.15 chiasmata and 8.62 chiasmata was observed respectively in *Tazip4-B2 ph1b* mutant-rye hybrids treated with water alone, and those treated with the Hoagland solution minus Mg^2+^ (Table 1). As expected, no significant differences were observed between the two treatments when double chiasmata were considered in these *Tazip4-B2 ph1b* hybrids.

Once the absence of Mg^2+^ was demonstrated to decrease homoeologous CO frequency in *Tazip4-B2 ph1b* mutant-rye hybrids, the effect of irrigating with only Mg^2+^ present at a final concentration of 1mM in water rather than in the Hoagland solution, was also analysed on homoeologous COs in *Tazip4-B2 ph1b* mutant-rye hybrids (Figure 1). Treatment with a solution containing only Mg^2+^ also increased homoeologous COs at metaphase I per meiocyte in these hybrids, showing no significant difference in comparison to the Hoagland solution treatment. A mean of 11.09 chiasmata was observed after treatment with 1mM Mg^2+^ in water, and 10.74 chiasmata after treatment with the Hoagland solution (Figure 1), when a single chiasma was considered. A similar situation was seen when double chiasmata were considered: no significant differences were observed in homoeologous COs per meiocyte in *Tazip4-B2 ph1b* mutant-rye hybrids after treatment with either 1mM Mg^2+^ or Hoagland solutions (Figure 1).

**FIGURE 1:**
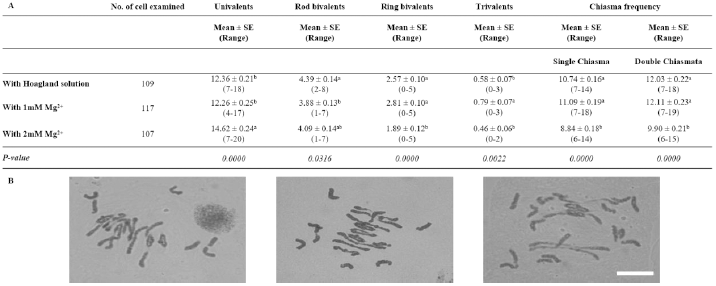
Effect of either 1mM or 2mM Mg^2+^ on homoeologous CO frequency of *T. aestivum* cv. Chinese Spring *Tazip4-B2 ph1b* mutant-rye hybrids. **(A)** Frequencies of univalents, bivalents, trivalents and chiasma frequency (single and double chiasmata) were scored at meiotic metaphase I in *Tazip4-B2 ph1b* mutant-rye hybrids treated with either Hoagland solution, 1mM Mg^2+^ or 2mM Mg^2+^ solution. Values in parenthesis indicate range of variation between cells. P < 0.05 indicates significant differences according to LSD test. **(B)** Representative meiotic configurations of *Tazip4-B2 ph1b* mutant-rye hybrids. From left to right: treatment with Hoagland solution, 1mM Mg^2+^ or 2mM Mg^2+^ solution. Bar: 20 μm.

The concentration of Mg^2+^ was subsequently increased to a final concentration of 2mM to assess whether the number of homoeologous COs could be increased further (Figure 1). Surprisingly, numbers of COs were reduced under these conditions (mean 11.09 for 1mM Mg^2+^ and 8.84 for 2mM Mg^2+^ treatments respectively, when single chiasma were considered, and 12.11 and 9.90, respectively, when double chiasmata were considered).

In addition to irrigating the plants with either Hoagland or Mg^2+^ solutions, we analysed the effect of treatment with 1mM Mg^2+^ in water following injection into the tillers of *Tazip4-B2 ph1b* mutant-rye hybrids. Injections were made just above each spike. Once again, homoeologous CO frequency was significantly increased in hybrids treated with 1mM Mg^2+^ when the solution was injected into the tiller (Table 2). A mean of 8.98 chiasmata in hybrids treated with water alone and 10.60 chiasmata in hybrids treated with 1mM Mg^2+^ was observed in the hybrids considering a single chiasma (Table 2) and a mean of 9.67 chiasmata and 11.30 chiasmata considering double chiasmata (Table 2).

**TABLE 2:** Effect of injecting 1mM Mg^2+^ solution into the tillers of *Tazip4-B2 ph1b* mutant-rye hybrids. Frequencies of univalents, bivalents, trivalents and chiasma frequency (single and double chiasmata) were scored at meiotic metaphase I in *Tazip4-B2 ph1b* mutant-rye hybrids treated with 1mM Mg^2+^ solution and with water alone. Values in parenthesis indicate range of variation between cells. P < 0.05 indicates significant differences according to LSD test.

|  | No. of cell<br>examined | Univalents | Rod bivalents | Ring bivalents | Trivalents | Chiasma frequency | Chiasma frequency |
| --- | --- | --- | --- | --- | --- | --- | --- |
|  |  | Mean ± SE<br>(Range) | Mean ± SE<br>(Range) | Mean ± SE<br>(Range) | Mean ± SE<br>(Range) | Mean ± SE<br>(Range) | Mean ± SE<br>(Range) |
|  |  |  |  |  |  | Single Chiasma | Double Chiasmata |
| Water alone | 87 | 15.02 ± 0.26<sup>a</sup><br>(3-20) | 3.43 ± 0.16<sup>b</sup><br>(0-7) | 2.12 ± 0.10<br>(0-4) | 0.63 ± 0.08<br>(0-3) | 8.98 ± 0.18<sup>b</sup><br>(6-16) | 9.67 ± 0.21<sup>b</sup><br>(6-17) |
| With 1mM Mg<sup>2+</sup> | 79 | 12.32 ± 0.26<sup>b</sup><br>(5-18) | 4.32 ± 0.17<sup>a</sup><br>(1-8) | 2.37 ± 0.12<br>(0-5) | 0.77 ± 0.08<br>(0-3) | 10.60 ± 0.19<sup>a</sup><br>(7-16) | 11.30 ± 0.22<sup>a</sup><br>(8-18) |
| *P-value* |  | 0.0000 | 0.0001 | 0.1079 | 0.2312 | 0.0000 | 0.0000 |

### Magnesium increases homoeologous COs in *Tazip4-B2 ph1b mutant*-*Ae. variabilis* hybrids

The addition of Mg^2+^ is thus identified as responsible for the increase in homoeologous CO at meiotic metaphase I in *Tazip4-B2 ph1b* mutant-rye hybrids. We then assessed the effect of 1mM Mg^2+^ on *T. aestivum* cv. Chinese spring *Tazip4-B2 ph1b* mutant-*Ae. variabilis* hybrids. Firstly, we scored the number of univalents, bivalents and multivalents, and total chiasma frequency in this hybrid, to compare the level of chiasma frequency to that previously reported by Kousaka and Endo, (2012) in *T. aestivum* cv. Chinese spring-*Ae. variabilis* hybrids in the absence of chromosome 5B. We observed a similar chiasma frequency in our hybrid (mean 14.15 chiasmata per meiocyte), to that previously reported in *T. aestivum* cv. Chinese spring-*Ae. variabilis* hybrids in the absence of chromosome 5B (mean of 14.09 chiasmata per meiocyte), confirming a similar level of meiotic metaphase I configuration in these hybrids.

We then analysed the effect of treatment with water alone and with either 1mM Mg^2+^ solution or complete Hoagland solution on the *Tazip4-B2 ph1b* mutant-*Ae. variabilis* hybrids. The total number of COs was significantly higher after treatment with 1mM Mg^2+^ than after treatment with water alone (without Mg^2+^ control), both in the case of single chiasma and double chiasmata, showing a mean of 15.31 and 14.15 chiasmata in the case of single chiasma, and a mean of 16.54 and 15.10 chiasmata in the case of double chiasmata, respectively (Figure 2). The number of univalents was significantly decreased and the number of trivalents was significantly increased when the plants were treated with 1mM Mg^2+^ solution in comparison to when Mg^2+^ was absent (a mean of 11.14 and 1.99, respectively, after treatment with 1mM Mg^2+^ and a mean of 12.49 and 1.60, respectively, after treatment with water alone were observed in Figure 2). With regard to the Hoagland solution treatment, significant differences were observed between hybrids treated with water alone and hybrids treated with Hoagland solution (Figure 2). Hoagland solution treatment showed the highest chiasma frequency, followed by 1mM Mg^2+^ and water alone (means of 16.53, 15.31 and 14.15 chiasmata were observed, respectively, when a single chiasma was considered, and means of 18.05, 16.54 and 15.10 chiasmata were observed, respectively, when double chiasmata were considered (Figure 2)).

**FIGURE 2:**
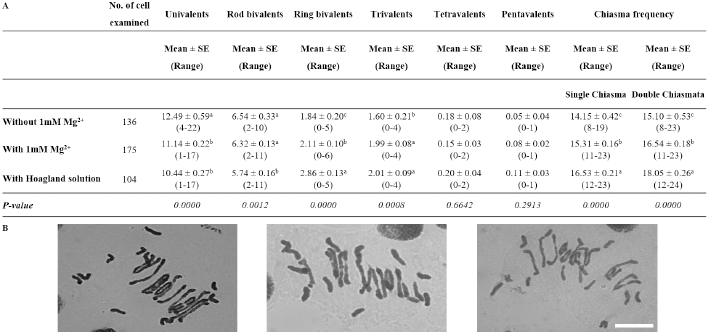
Effect of either 1mM Mg^2+^ or Hoagland solution on homoeologous CO frequency of *T. aestivum* cv. Chinese Spring *Tazip4-B2 ph1b* mutant-*Ae. variabilis* hybrids. **(A)** Frequencies of univalents, bivalents, trivalents, tetravalents, pentavalents and chiasma frequency (single and double chiasmata) were scored at meiotic metaphase I in *Tazip4-B2 ph1b* mutant-*Ae. variabilis* hybrids treated with either 1mM Mg^2+^ or Hoagland solution. Values in parenthesis indicate range of variation between cells. P < 0.05 indicates significant differences according to LSD test. **(B)** Representative meiotic configurations of *Tazip4-B2 ph1b* mutant-*Ae. variabilis* hybrids. From left to right: water alone, 1mM Mg^2+^ treatment and Hoagland solution treatment. Bar: 20 μm.

**FIGURE 3:**
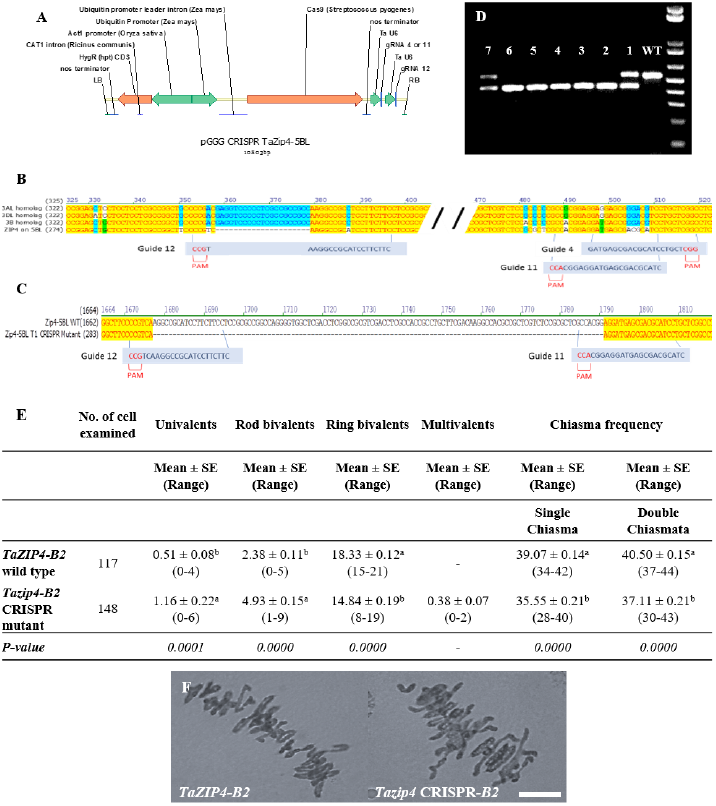
Development and phenotypic analysis of *Tazip4-B2* CRISPR mutants generated using RNA-guided Cas9. **(A)** Schematic of the structure of the pGGG CRISPR *TaZip4-B2* vector used in this study. **(B)** Alignment of all copies of the *ZIP4* gene in wheat showing sequences and positions of the three sgRNAs designed to specifically target *TaZIP4-B2* with their corresponding protospacer-adjacent motif (PAM). **(C)** Alignment of *TaZIP4-B2* wild type and *Tazip4-B2 CRISPR* mutant sequences sowing the localization of the large deletion (115 bp) in *TaZIP4-B2*. **(D)** Genotypic assays for the identification of homozygous edited lines (lines: #2, #3, #4, #5 and #6) and heterozygous lines (lines: #1 and #7). Wild-type Fielder (WT) was used as a control line. **(E)** Frequencies of univalents, bivalents and multivalents, and total chiasma frequency (single and double chiasmata) were scored at meiotic metaphase I in wild-type Fielder and *Tazip4-B2* CRISPR mutant. Values in parenthesis indicate range of variation between cells. P < 0.05 indicates significant differences according to Dunn’s test. **(F)** Representative meiotic metaphase I configurations of wild-type Fielder and *Tazip4-B2* CRISPR Fielder mutants. Left: wheat cv. Fielder and right: *Tazip4-B2* CRISPR mutant. Bar: 20 μm.

### Magnesium increases homoeologous COs in wheat *Tazip4-B2* TILLING mutant-*Ae variabilis* mutant hybrids

Recently we reported that *Tazip4-B2* TILLING mutants crossed with *Ae. variabilis* exhibited homoeologous COs at meiotic metaphase I. We therefore decided to analyse whether the level of homoeologous COs induced by *Tazip4-B2* TILLING mutants was also affected by treatment with 1mM Mg^2+^ solution. To assess the effect of 1mM Mg^2+^ on homoeologous CO frequency at metaphase I, we added 100 mL per plant of a solution of either 1mM Mg^2+^ in water or Hoagland solution once a week to the soil in which these hybrids were growing. In this experiment, we analysed both *Tazip4-B2* TILLING mutant lines (Cad1691 and Cad0348) (Rey et al., 2017), crossed with *Ae. variabilis*. Both TILLING mutant hybrids showed a significant increase in chiasma frequency after treatment with 1mM Mg^2+^, compared to chiasma frequency obtained in both the hybrids treated with water alone. The *Tazip4-B2* TILLING mutant (Cad1691)-*Ae. variabilis* and the *Tazip4-B2* TILLING mutant (Cad0348)-*Ae. variabilis* hybrids showed means of 13.41 and 13.66 single chiasma frequency, respectively, after treatment with 1mM Mg^2+^ and means of 12.21 and 12.23 single chiasma frequency, respectively, in water alone (Table 3; Figure S2). Significant differences were also observed when double chiasmata were scored in both mutant lines (Table 3; Figure S2). Numbers of univalents and trivalents were also affected by treatment with 1mM Mg^2+^ in both mutant lines as in the *Tazip4-B2 ph1b* mutant-*Ae. variabilis* hybrids described in the previous section. Numbers of univalents were significantly decreased both in *Tazip4-B2* TILLING mutant (Cad1691)-*Ae. variabilis* and *Tazip4-B2* TILLING mutant (Cad0348)-*Ae. variabilis* hybrids treated with 1mM Mg^2+^ (means of 12.74 and 12.11 univalents respectively with Mg^2+^ and means of 14.74 and 14.63 univalents respectively with water alone (Table 3)). Numbers of trivalents were significantly increased both in wheat (Cad1691)-*Ae. variabilis* and wheat (Cad0348)-*Ae. variabilis* hybrids, after treatment with 1mM Mg^2+^ (means of 1.63 and 1.93 trivalents respectively with Mg^2+^, and means of 1.05 and 1.27 trivalents respectively with water alone (Table 3)).

**TABLE 3:** Effect of either 1mM Mg^2+^ or Hoagland solution on the homoeologous CO frequency of *T. aestivum* cv. Cadenza (Cad1691-*Tazip4-B2*)-*Ae. variabilis* and *T. aestivum* cv. Cadenza (Cad0348-*Tazip4-B2*)-*Ae. variabilis* hybrids. Frequencies of univalents, bivalents, trivalents, tetravalents, pentavalents and chiasma frequency (single and double chiasmata) were scored at meiotic metaphase I in *Tazip4-B2* TILLING mutant-*Ae. variabilis* hybrids treated with either 1mM Mg^2+^ or Hoagland solution. Values in parenthesis indicate range of variation between cells. P < 0.05 indicates significant differences according to LSD test. ^*^This data published in Rey et al., (2017).

|  |  | No. of cell<br>examined | Univalents | Rod bivalents | Ring bivalents | Trivalents | Tetravalents | Pentavalents | Chiasma frequency |  |
| --- | --- | --- | --- | --- | --- | --- | --- | --- | --- | --- |
|  |  |  | Mean ± SE<br>(Range) | Mean ± SE<br>(Range) | Mean ± SE<br>(Range) | Mean ± SE<br>(Range) | Mean ± SE<br>(Range) | Mean ± SE<br>(Range) | Mean ± SE<br>(Range) | Mean ± SE<br>(Range) |
|  |  |  |  |  |  |  |  |  | Single Chiasma | Double Chiasmata |
| Water alone* | Cad1691 x *Ae. variabilis* hybrids | 106 | 14.74 ± 0.29<sup>a</sup><br>(7-26) | 6.75 ± 0.17<br>(3-11) | 1.26 ± 0.08<br>(0-4) | 1.05 ± 0.08<sup>b</sup><br>(0-4) | 0.22 ± 0.04<br>(0-2) | 0.03 ± 0.02<br>(0-1) | 12.21 ± 0.19<sup>b</sup><br>(8-18) | 12.74 ± 0.21<sup>b</sup><br>(8-20) |
|  | Cad0348 x *Ae. variabilis* hybrids | 102 | 14.63 ± 0.28<sup>a</sup><br>(6-21) | 6.64 ± 0.18<br>(3-10) | 1.12 ± 0.10<br>(0-4) | 1.27 ± 0.11<sup>b</sup><br>(0-4) | 0.21 ± 0.04<br>(0-1) | 0.05 ± 0.02<br>(0-1) | 12.23 ± 0.20<sup>b</sup><br>(7-19) | 12.68 ± 0.23<sup>b</sup><br>(7-21) |
| With 1mM<br>Mg<sup>2+</sup> | Cad1691 x *Ae. variabilis* hybrids | 95 | 12.74 ± 0.33<sup>b</sup><br>(5-19) | 7.06 ± 0.20<br>(2-11) | 1.18 ± 0.11<br>(0-3) | 1.63 ± 0.13<sup>a</sup><br>(0-5) | 0.17 ± 0.30<br>(0-1) | 0.04 ± 0.02<br>(0-1) | 13.41 ± 0.22<sup>a</sup><br>(8-18) | 13.75 ± 0.23<sup>a</sup><br>(8-19) |
|  | Cad0348 x *Ae. variabilis* hybrids | 110 | 12.11 ± 0.32<sup>b</sup><br>(3-20) | 6.93 ± 0.18<br>(1-12) | 1.51 ± 0.11<br>(0-4) | 1.93 ± 0.11<sup>a</sup><br>(0-5) | 0.24 ± 0.04<br>(0-2) | 0.07 ± 0.02<br>(0-1) | 14.21 ± 0.23<sup>a</sup><br>(9-21) | 14.90 ± 0.25<sup>a</sup><br>(10-22) |
| With Hoagland<br>solution | Cad1691 x *Ae. variabilis* hybrids | 118 | 11.94 ± 0.33<sup>b</sup><br>(2-20) | 6.76 ± 0.23<br>(0-12) | 1.17 ± 0.10<br>(0-4) | 1.50 ± 0.11<sup>a</sup><br>(0-5) | 0.26 ± 0.05<br>(0-2) | 0.04 ± 0.02<br>(0-1) | 13.66 ± 0.21<sup>a</sup><br>(9-20) | 14.14 ± 0.22<sup>a</sup><br>(9-21) |
|  | Cad0348 x *Ae. variabilis* hybrids | 142 | 12.48 ± 0.25<sup>b</sup><br>(2-20) | 6.70 ± 0.17<br>(1-12) | 1.37 ± 0.09<br>(0-4) | 1.76 ± 0.10<sup>a</sup><br>(0-5) | 0.23 ± 0.04<br>(0-2) | 0.04 ± 0.02<br>(0-1) | 13.84 ± 0.19<sup>a</sup><br>(9-21) | 14.30 ± 0.21<sup>a</sup><br>(9-22) |
| P-value | Cad1691 x *Ae. variabilis* hybrids |  | 0.0000 | 0.1930 | 0.6630 | 0.0026 | 0.3278 | 0.9376 | 0.0000 | 0.0000 |
|  | Cad0348 x *Ae. variabilis* hybrids |  | 0.0000 | 0.3220 | 0.5967 | 0.0003 | 0.9720 | 0.3650 | 0.0000 | 0.0000 |

Finally, we assessed the effect of treating with Hoagland solution and with water alone, finding significant differences in homoeologous COs between the two treatments, both in *Tazip4-B2* TILLING mutant (Cad1691)-*Ae. variabilis* and *Tazip4-B2* TILLING mutant (Cad0348)-*Ae. variabilis* hybrids. Numbers of univalents and trivalents were also affected to the same extent (Table 3). In the *Tazip4-B2* TILLING mutant (Cad1691)-*Ae. variabilis* hybrid, means of 14.74 univalents and 1.05 trivalents were observed in hybrids treated with water alone, and means of 11.94 univalents and 1.50 trivalents observed in hybrids treated with Hoagland solution (Table 3). In the *Tazip4-B2* TILLING mutant (Cad0348)-*Ae. variabilis* hybrid, means of 14.63 univalents and 1.27 trivalents were observed in hybrids with water alone and means of 12.48 univalents and 1.76 trivalents in hybrids treated with Hoagland solution (Table 3).

### Phenotypic analysis of *Tazip4-B2* mutants generated by CRISPR/Cas9 system

Firstly, eighty-one primary transgenic plants (T0) were analysed by PCR followed by direct sequencing. Four plants were identified with edits in the target region. One plant had a perfect 115bp deletion between guides G11 and G12. Twenty-four T1 plants from this line were screened and 5 homozygous edited plants with the 115bp deletion were recovered. These plants were used to score the number of univalents, bivalents and multivalents, and total chiasma frequency in the *Tazip4-B2* mutant CRISPR lines (Figure 3). Wild-type Fielder lines were used as control plants (Figure 3). The *Tazip4-B2* CRISPR mutant lines exhibited a significant reduction in ring bivalents, from a mean of 18.33 to 14.84 in the wild-type Fielder and CRISPR mutant lines respectively (Figure 3). A significant increase in the number of univalents and rod bivalents was also observed, from means of 0.51 univalents and 2.38 rod bivalents in the wild-type Fielder line, to means of 1.16 univalents and 4.93 rod bivalents in the CRISPR mutant lines (Figure 3). This indicates a significant reduction in homologous COs in these *Tazip4-B2* mutant lines (Figure 3). Chiasma frequency decreased from a mean of 39.07 single chiasma and 40.50 double chiasmata in the wild-type Fielder line, to a mean of 35.55 single chiasma and 37.11 double chiasmata in the *Tazip4-B2* CRISPR mutant (Figure 3).

### Magnesium also increases homoeologous COs in wheat *Tazip4-B2* CRISPR mutant-*Ae. variabilis* mutant hybrids

For this study, a wild-type Fielder and a *Tazip4-B2* CRISPR Fielder mutant line were crossed with *Ae. variabilis* to assess the level of homoeologous COs in the resulting hybrids (Table S2). Frequency of univalents, bivalents and multivalents, and total chiasma frequency were scored at meiotic metaphase I (Table S2). *Tazip4-B2* CRISPR mutant hybrids exhibited a significant increase in single chiasma frequency, from a mean of 3.15 in the wild-type Fielder-*Ae.variabilis* hybrid to 16.66 in the *Tazip4-B2* CRISPR-*Ae. variabilis* hybrid (Table S2). Double chiasma frequency was also increased in the *Tazip4-B2* CRISPR mutant hybrids (Table S2). There was also a similar increase in the chiasma frequency to that reported previously in *Tazip4-B2* TILLING-*Ae. variabilis* hybrids (Rey et al., (2017)).

Having observed the effect of treatment with Mg^2+^ on homoeologous CO frequency in the *Tazip4-B2* TILLING mutant hybrids, we also analysed the effect of this ion on *Tazip4-B2* CRISPR mutants-*Ae. variabilis* hybrids. We added 100 mL of a solution of 1mM Mg^2+^ in water or Hoagland solution once a week to the soil in which the hybrids were growing. As expected, the addition of nutrients to these mutant hybrids caused a significant increase in chiasma frequency (Table 4). *Tazip4-B2* CRISPR-*Ae. variabilis* hybrids treated with water alone exhibited means of 16.66 single chiasma frequency and 18.10 double chiasma frequency. Addition of 1mM Mg^2+^ caused a significant increase in chiasma frequency of these mutant hybrids (means of 17.67 and 18.75 single and double chiasma frequency respectively) (Table 4). Also, the addition of Hoagland solution increased the homoeologous COs in these *Tazip4-B2* CRISPR hybrids. Means of 18.34 and 19.82 single and double chiasma frequency respectively, were observed in those plants treated with Hoagland solution (Table 4).

**TABLE 4:** Effect of either 1mM Mg^2+^or Hoagland solution on the homoeologous CO frequency of wheat *Tazip4-B2* CRISPR-*Ae. variabilis* mutant hybrids. Frequencies of univalents, bivalents, trivalents, tetravalents, pentavalents and chiasma frequency (single and double chiasmata) were scored at meiotic metaphase I in *Tazip4-B2* CRISPR mutant-*Ae. variabilis* hybrids treated with either 1mM Mg^2+^ or Hoagland solution. Values in parenthesis indicate range of variation between cells. P < 0.05 indicates significant differences according to LSD test.

|  | No. of cell<br>examined | Univalents | Rod bivalents | Ring bivalents | Trivalents | Tetravalents | Pentavalents | Hexavalents | Chiasma frequency |  |
| --- | --- | --- | --- | --- | --- | --- | --- | --- | --- | --- |
|  |  | Mean ± SE<br>(Range) | Mean ± SE<br>(Range) | Mean ± SE<br>(Range) | Mean ± SE<br>(Range) | Mean ± SE<br>(Range) | Mean ± SE<br>(Range) | Mean ± SE<br>(Range) | Mean ± SE<br>(Range) | Mean ± SE<br>(Range) |
|  |  |  |  |  |  |  |  |  | Single Chiasma | Double Chiasmata |
| Water alone | 124 | 9.64 ± 0.27<sup>a</sup><br>(3-17) | 5.64 ± 0.17<sup>b</sup><br>(2-10) | 1.94 ± 0.11<sup>b</sup><br>(0-6) | 2.37 ± 0.11<br>(0-6) | 0.52 ± 0.06<br>(0-3) | 0.20 ± 0.04<sup>a</sup><br>(0-2) | 0.00 ± 0.00<br>(0-0) | 16.66 ± 0.21<sup>c</sup><br>(11-22) | 18.10 ± 0.23<sup>c</sup><br>(12-24) |
| With 1mM Mg<sup>2+</sup> | 135 | 7.92 ± 0.25<sup>b</sup><br>(0-14) | 6.19 ± 0.15<sup>a</sup><br>(3-11) | 2.02 ± 0.09<sup>b</sup><br>(0-6) | 2.68 ± 0.09<br>(0-3) | 0.50 ± 0.06<br>(0-2) | 0.10 ± 0.01<sup>b</sup><br>(0-1) | 0.01 ± 0.01<br>(0-1) | 17.67 ± 0.17<sup>b</sup><br>(14-23) | 18.75 ± 0.20<sup>b</sup><br>(14-24) |
| With Hoagland<br>solution | 125 | 7.90 ± 0.28<sup>b</sup><br>(0-15) | 5.56 ± 0.17<sup>b</sup><br>(1-11) | 2.82 ± 0.11<sup>a</sup><br>(0-6) | 2.53 ± 0.11<br>(0-5) | 0.59 ± 0.07<br>(0-3) | 0.08 ± 0.03<sup>b</sup><br>(0-2) | 0.00 ± 0.00<br>(0-0) | 18.34 ± 0.21<sup>a</sup><br>(14-24) | 19.82 ± 0.23<sup>a</sup><br>(15-25) |
| P-value |  | 0.0000 | 0.0121 | 0.0000 | 0.1021 | 0.5929 | 0.0259 | 0.1575 | 0.0000 | 0.0000 |

## Discussion

Introgression of genetic material from relative species into bread wheat has been used in plant breeding for over 50 years, although classical plant breeding methods to introgress wild relative segments into wheat are both inefficient and time consuming (Ko et al., 2002). Recent availability of SNP based arrays, combined with classical cytogenetic approaches, significantly enhanced our ability to exploit wild relatives (King et al., 2017a, 2017b), using lines carrying a deletion of either the whole of chromosome 5B, or a smaller 70Mb segment (*ph1b*) (Riley and Chapman, 1958; Sears and Okamoto, 1958: Sears, 1977), to increase the level of homoeologous crossovers between wild relatives and wheat chromosomes. Recombination between wild relative chromosomes and wheat chromosomes is, however, still limited. Thus, there is a need to find abiotic or biotic treatments such as temperature, nutritional availability, DNA-damaging agents, among others (Lambing et al., 2017) to enhance recombination. Martín et al., (2017) recently reported an alternative tool to increase CO number in *Tazip4-B2 ph1b* mutant-rye hybrids, using the addition of a Hoagland solution to the soil in which the plants are grown. Martín et al., (2017) also showed that the presence of the Hoagland solution did not affect the homoeologous CO number in wild-type wheat-rye hybrids.

Here, we report the successful identification of the particular Hoagland solution constituent responsible for the observed increase in homoeologous CO frequency. After analysing *Tazip4-B2 ph1b* mutant-rye hybrids in the absence of each separate Hoagland solution macronutrient, we observed a significant reduction in homoeologous CO frequency when the Mg^2+^ ion was absent. This suggests that the Mg^2+^ ion is mainly responsible for the effect of Hoagland solution on homoeologous COs described previously by Martín et al., (2017). These observations were obtained after cytogenetic analysis of meiotic configurations at meiotic metaphase I. The analysis involved scoring single and double chiasmata in the chromosomal structures (Figure S1). Single chiasma counting has commonly been used in many studies to measure chiasma frequency in wheat (Sears, 1977; Dhaliwal et al., 1977; Roberts et al., 1999). However, other studies have suggested that double chiasmata may occur in these chromosomal configurations (Gennaro et al., 2012; Dreissig et al., 2017). Double chiasmata were considered in the present study, as a high number of MLH1 sites were previously reported in *Tazip4-B2 ph1b* mutant-rye hybrids in Martín et al., (2014). In our studies, up to 19 chiasmata were scored in *Tazip4-B2 ph1b* mutant-rye hybrids, which is similar to the number of MLH1 sites observed previously (Martín et al., 2014).

The effect of treatment with a solution of 1mM Mg^2+^ in water, was analysed to confirm whether that the Mg^2+^ ion was responsible for the increase in homoeologous COs observed in these hybrids. The effect of treatment with this solution was assessed either by irrigation of, or injection into *Tazip4-B2 ph1b* mutant-rye hybrids. Surprisingly, both methods of application increased homoeologous CO frequency in the *Tazip4-B2 ph1b* mutant-rye hybrids. Thus, the results from the injection method of application suggested that the 1mM Mg^2+^ concentration was directly responsible for the increased homoeologous CO effect seen in the *Tazip4-B2 ph1b* mutant-rye hybrids, rather than through indirect effects on the plant growth or development. However, homoeologous CO frequency was decreased when the Mg^2+^ concentration was increased further (Figure 1). This reduction in COs was associated with a significant increase in the number of univalents, and decrease in the number of ring bivalents and trivalents.

A recent study revealed that *TaZIP4-B2* within the 5B region defined by the 70Mb *ph1b* deletion, was responsible for the suppression of homoeologous COs in hybrids (Rey et al., 2017). *Tazip4-B2* TILLING mutants (one with a missense mutation and another with a nonsense mutation), when crossed with *Ae. variabilis*, exhibit similar levels of homoeologous CO that observed in *ph1b-Ae. variabilis* hybrids (Rey et al., 2017). It was therefore important to assess the effect of 1mM Mg^2+^ solution on these *Tazip4-B2* TILLING mutant-*Ae. variabilis* hybrids to confirm that the effect was associated with *Tazip4-B2*, and that the Mg^2+^ effect could also be observed in a different hybrid. Moreover, we also applied the CRISPR/Cas9 genome editing system in hexaploid wheat cv. Fielder to the mutant *TaZIP4-B2* to compare its mutant phenotype with those observed in TILLING mutant lines, and their *Ae. variabilis* hybrids. *Tazip4-B2* CRISPR mutants showed a significant decrease in homologous COs compared to control plants (*TaZIP4-B2* wild type wheat), similar to that already reported for *Tazip4-B2* TILLING mutants (Rey et al., 2017). Also, as expected, a significant increase was observed in *Tazip4-B2* CRISPR mutant-*Ae. variabilis* hybrids, similar to that observed in both *ph1b*-*Ae. variabilis* and *Tazip4-B2* TILLING mutant-*Ae. variabilis* hybrids. Furthermore, the addition of 1mM Mg^2+^ to all these hybrids increased the frequency of homoeologous CO. This confirms that the Mg^2+^ effect is associated with *Tazip4-B2*, and occurs in different hybrids. The only difference observed with the *Tazip4-B2* CRISPR and TILLING mutants was the occurrence of multivalents in the CRISPR mutants compared to the TILLING mutants (Figure 3 and Rey et al., 2017). This suggests that *TaZIP4-B2* not only promotes homologous COs and restricts homoeologous COs, but also contributes to the efficiency of homologous pairing. We hypothesize that the CRISPR deletion disrupts more of the *TaZIP4-B2* function than the TILLING mutants. Interestingly, in rice, *ZIP4* mutants have previously been reported to show a delay in completing homologous synapsis (Shen et al., 2012), however, in that diploid species, this does not lead to homoeologous COs because only homologues are present. However, in the *ph1b* mutant, delayed pairing of some homologues is observed until after the telomere bouquet, allowing some subsequent homoeologous pairing to take place. This delayed pairing of homologues in the *ph1b* mutant is consistent with a *ZIP4* mutant phenotype.

Magnesium is one of the most important nutrients, mainly involved in the general promotion of plant growth and development. In terms of CO function, Mg^2+^ may affect multiple proteins in the class I interference crossover pathway either in a positive or negative manner. For example, recent studies have suggested that Mg^2+^ is required for the endonuclease activity of the MLH1-MLH3 heterodimer (Rogacheva et al., 2014). The MLH1-MLH3 heterodimer shows a strong preference for HJs in the absence of Mg^2+^ (Ranjha et al., 2014). Whatever the target, our present study reveals that homoeologous COs can be increased by the 1mM Mg^2+^ treatment of *Tazip4-B2* (*ph1b*, TILLING or CRISPR derived) mutant-wild relative hybrids. Thus, this treatment can be used as a tool to enhance the introgression of wild relative traits into wheat.

## Author Contributions

M-DR, AM, PS and GM conceived and designed the study. MS, SH and WH participated in the development of the *Tazip4-B2* mutant in bread wheat cv. Fielder by CRISPR/Cas9 system. M-DR analysed the research results and wrote the first draft. PS and GM modified the paper. All authors have read and approved the final version of the manuscript.

## Acknowledgements

The authors thank Ali Pendle (John Innes Centre, UK) for her valuable comments in the writing of the manuscript. This work was supported by the UK Biotechnology and Biological Research Council (BBSRC), through a grant part of the Designing Future Wheat (DFW) Institute Strategic Programme (BB/P016855/1), three grants (Grant BB/J004588/1; Grant BB/M009599/1; Grant BB/J007188/1); and by a Marie Curie Fellowship Grant (H2020-MSCA-IF-2015-703117).

## Supporting information

**TABLE S1:** Genotypes and number of plants used for analysing the effect of a nutrient solution in homoeologous CO frequency in wheat and its relative species.

| Genotype | Treatment | No. of plants |
| --- | --- | --- |
| **Absence of the *Ph1* locus** |  |  |
| CS- x Rye hybrids | Hoagland Solution | 5 |
| CS- x Rye hybrids | without Hoagland | 5 |
| CS- x Rye hybrids | with Hoagland Solution - NH<sub>2</sub> H<sub>2</sub>PO<sub>4</sub> | 5 |
| CS- x Rye hybrids | with Hoagland Solution - KNO<sub>3</sub> | 5 |
| CS- x Rye hybrids | with Hoagland Solution - CaNO<sub>3</sub> | 5 |
| CS- x Rye hybrids | with Hoagland Solution - MgSO<sub>4</sub> | 5 |
| CS- x Rye hybrids | with Hoagland | 5 |
| CS- x Rye hybrids | 1mM Magnesium | 5 |
| CS- x Rye hybrids | 2mM Magnesium | 5 |
| CS- x *Ae. variabilis* hybrids | without 1mM Magnesium | 4 |
| CS- x *Ae. variabilis* hybrids | 1mM Magnesium | 5 |
| CS- x *Ae. variabilis* hybrids | Hoagland Solution | 3 |
| **Absence of the *TaZIP4* gene** |  |  |
| **TILLING** |  |  |
| Cad1691 x *Ae. variabilis* hybrids | without 1mM Magnesium | 5 |
| Cad1691 x *Ae. variabilis* hybrids | 1mM Magnesium | 4 |
| Cad1691 x *Ae. variabilis* hybrids | Hoagland Solution | 4 |
| Cad0348 x *Ae. variabilis* hybrids | without 1mM Magnesium | 5 |
| Cad0348 x *Ae. variabilis* hybrids | 1mM Magnesium | 4 |
| Cad0348 x *Ae. variabilis* hybrids | Hoagland Solution | 4 |
| **CRISPR/Cas9 system** |  |  |
| Wheat cv. Fielder carrying *TaZIP4-B2* |  | 5 |
| Wheat cv. Fielder lacking *TaZIP4-B2* |  | 5 |
| *TaZIP4-B2* - *Ae. variabilis* hybrids |  | 4 |
| *Tazip4-B2* CRISPR - *Ae. variabilis* hybrids | without 1mM Magnesium | 4 |
| *Tazip4-B2* CRISPR - *Ae. variabilis* hybrids | 1mM Magnesium | 4 |
| *Tazip4-B2* CRISPR - *Ae. variabilis* hybrids | Hoagland Solution | 4 |

**TABLE S2:** Frequencies of univalents, bivalents, multivalents and chiasma frequency (single and double chiasmata) were scored at meiotic metaphase I in wheat *Tazip4-B2* CRISPR mutant *Ae. variabilis* hybrids. Values in parenthesis indicate range of variation between cells. P < 0.05 indicates significant differences according to Dunn’s test.

|  | No. of cell<br>examined | Univalents | Rod bivalents | Ring<br>bivalents | Trivalents | Tetraivalents | Pentavalents | Chiasma frequency |  |
| --- | --- | --- | --- | --- | --- | --- | --- | --- | --- |
|  |  | Mean ± SE<br>(Range) | Mean ± SE<br>(Range) | Mean ± SE<br>(Range) | Mean ± SE<br>(Range) | Mean ± SE<br>(Range) | Mean ± SE<br>(Range) | Mean ± SE<br>(Range) | Mean ± SE<br>(Range) |
|  |  |  |  |  |  |  |  | Single Chiasma | Double Chiasmata |
| Fielder x *Ae. variabilis* hybrids | 172 | 28.99 ± 0.27<sup>a</sup><br>(20-35) | 2.61 ± 0.12<sup>b</sup><br>(0-7) | 0.05 ± 0.02<sup>b</sup><br>(0-1) | 0.23 ± 0.04<sup>b</sup><br>(0-2) | - | - | 3.15 ± 0.15<sup>b</sup><br>(0-8) | 3.41 ± 0.17<sup>b</sup><br>(0-7) |
| CRISPR x *Ae. variabilis* hybrids | 124 | 9.64 ± 0.27<sup>b</sup><br>(3-17) | 5.64 ± 0.17<sup>a</sup><br>(2-10) | 1.94 ± 0.11<sup>a</sup><br>(0-6) | 2.37 ± 0.11<sup>a</sup><br>(0-6) | 0.52 ± 0.06<br>(0-3) | 0.20 ± 0.04<br>(0-2) | 16.66 ± 0.21<sup>a</sup><br>(11-22) | 18.10 ± 0.23<sup>a</sup><br>(12-24) |
| *P-value* |  | 0.0000 | 0.0000 | 0.0000 | 0.0000 | - | - | 0.0000 | 0.0000 |

**FIGURE S1:**
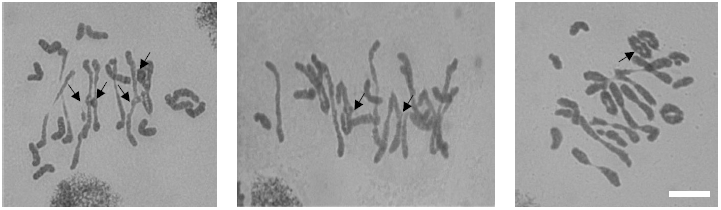
Chromosomal configurations with single chiasma or double chiasmata highlighted with arrows. These structures marked by an arrow were counted as either single or double chiasmata in all analysed meiocytes. Both datasets are shown in all analysed genotypes. Bar: 20 μm.

**FIGURE S2:**
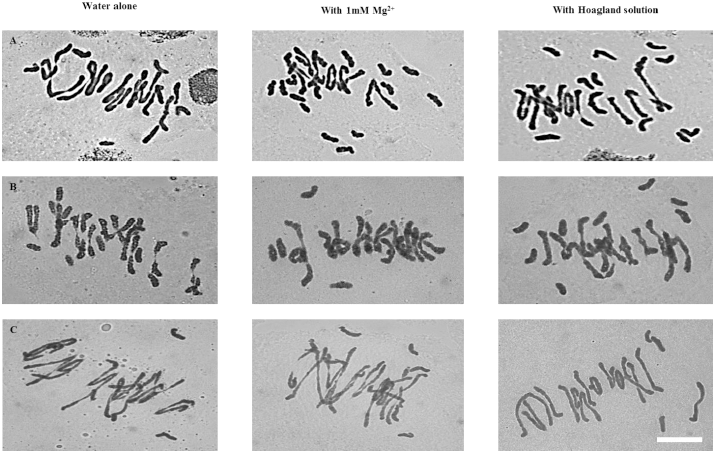
Representative meiotic configurations of *Triticum aestivum* cv. Cadenza (Cad1691-*Tazip4-B2* mutant)-*Ae. variabilis* (A) and *Triticum aestivum* cv. Cadenza (Cad0348-*Tazip4-B2* mutant)-*Ae. Variabilis* (B) and wheat *Tazip4-B2* CRISPR mutant-*Ae. variabilis* mutant (c) hybrids. From left to right: water alone, treated with either 1mM Mg^2+^ or Hoagland solution. Bar: 20 μm.

